# Dissociable contributions of the medial parietal cortex to recognition memory

**DOI:** 10.1101/2023.09.12.557048

**Authors:** Seth R. Koslov, Joseph W. Kable, Brett L. Foster

## Abstract

Human neuroimaging studies of episodic memory retrieval routinely observe the engagement of specific cortical regions beyond the medial temporal lobe. Of these, medial parietal cortex (MPC) is of particular interest given its ubiquitous, and yet distinct, functional characteristics during different types of retrieval tasks. Specifically, while recognition memory and autobiographical recall tasks are both used to probe episodic retrieval, these paradigms consistently drive distinct patterns of response within MPC. This dissociation adds to growing evidence suggesting a common principle of functional organization across memory related brain structures, specifically regarding the control or content demands of memory-based decisions. To carefully examine this putative organization, we used a high-resolution fMRI dataset collected at ultra-high field (7T) while subjects performed thousands of recognition-memory trials to identify MPC regions responsive to recognition-decisions or semantic content of stimuli within and across individuals. We observed interleaving, though distinct, functional subregions of MPC where responses were sensitive to either recognition decisions or the semantic representation of stimuli, but rarely both. In addition, this functional dissociation within MPC was further accentuated by distinct profiles of connectivity bias with the hippocampus during task and rest. Finally, we show that recent observations of person and place selectivity within MPC reflect category specific responses from within identified semantic regions that are sensitive to mnemonic demands. Together, these data better account for how distinct patterns of MPC responses can occur as a result of task demands during episodic retrieval and may reflect a common principle of organization throughout hippocampal-neocortical memory systems.

**Significance statement:** Medial parietal cortex (MPC) plays a growing role in contemporary theories of episodic memory, as it is reliably observed in human neuroimaging to be engaged during tasks of recognition and retrieval. However, the spatial pattern of MPC engagement consistently differs across these putatively similar episodic memory tasks. Despite a large literature indicating that the MPC is important for episodic memory, there is little consensus about its specific role. Here, we employed ‘precision-neuroimaging’ to identify dissociable interleaving MPC subregions, where activity reflected either memory-based decision-making or stimulus content. This dissociation within MPC provides a better understanding for how retrieval demands shape response patterns and speaks to growing evidence for a common principle of organization across memory structures of the human brain.

## Introduction

An essential feature of episodic memory is the ability to retrieve, reconstruct, or recognize prior experiences. While these cognitive abilities are classically associated with the medial temporal lobe (MTL) and hippocampus^2,3^, contemporary theories of episodic memory also emphasize the critical involvement of specific cortical regions^4–6^. Of these regions, the medial parietal cortex (MPC; Figure 1a) is of particular interest given that it is routinely observed in neuroimaging studies of episodic memory retrieval^7–12^, demonstrates larger magnitude responses with increasing memory strength^13,14^, and displays strong anatomical connectivity with MTL structures^15,16^. However, despite the ubiquitous observation of MPC activity during episodic retrieval, little is known about this region’s unique contributions to mnemonic processing and therefore its role in cognition more broadly^17^. Traditionally, investigators have used tasks requiring the recall of autobiographical events, or more commonly, laboratory-based tasks of studied item recognition, to experimentally probe episodic memory^18,19^. Strikingly, it has long been observed that despite both kinds of task being viewed as assays for episodic retrieval, the MPC displays distinct patterns of response to each of these tasks^20,21^ (see Figure 1b). Gaining a better understanding of this marked functional dissociation is important for not only characterizing the role of the MPC in mnemonic processes, but also for better understanding the mechanisms that support distinct cognitive stages and demands during recall and recognition.

**Figure 1.**
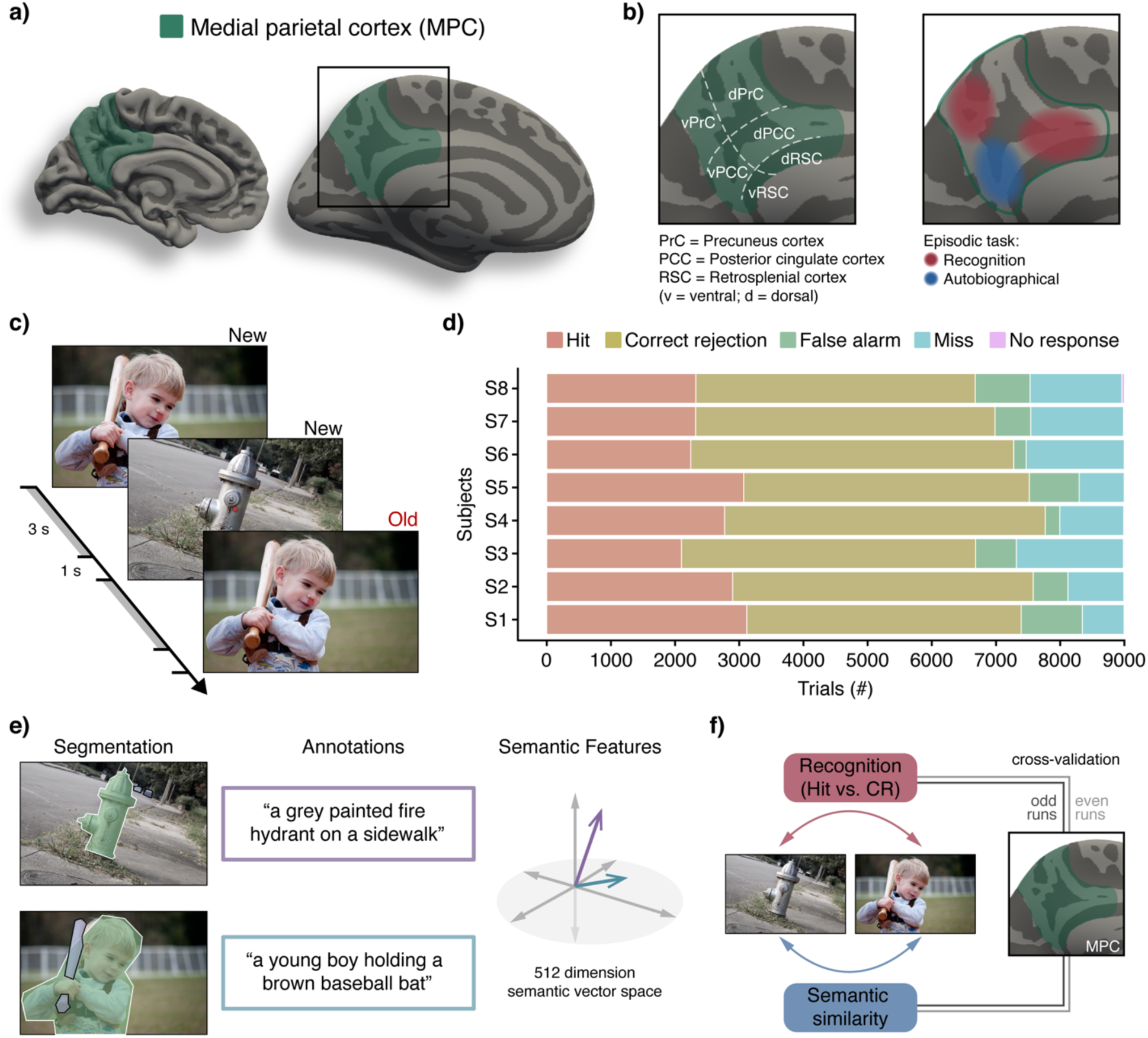
MPC anatomy, NSD experimental paradigm and analysis approach. **a**) MPC reflects the medial aspect of the parietal lobe, also termed posteromedial cortex (PMC). Demarcation of the MPC (green) is show on a standard (left) and inflated (right) cortical surface. **b)** MPC contains clear cytoarchitectural subdivisions, comprised of the precuneus (PrC), posterior-cingulate (PCC) and retrosplenial (RSC) cortices (left). Schematic (right) depicts common dissociable spatial patterns observed within MPC by fMRI studies of recognition (red) and autobiographical (blue) tasks of episodic retrieval. **c)** During the NSD stimulus-recognition task, subjects were shown one image at a time (3s) and asked to indicate whether the image was ‘old’ or ‘new’, followed by a brief fixation dot (1s). **d)** The NSD includes data from 8 subjects as they performed the stimulus-recognition task over the course of multiple sessions. Our analyses focus on a subset of those sessions (sessions 1-12), which included memory-decisions spanning up to 100 days since the time of encoding (9,000 trials). **e)** Stimuli are from the COCO image dataset, which includes detailed image content segmentation and associated text annotations. These text annotations were used to perform semantic similarity analysis across stimuli (see Materials and Methods). **f)** Data analyses focused on identifying MPC responses sensitive to the mnemonic status (recognition regions) and semantic content (semantic regions) of presented stimuli. Given the focus on single-subject level analysis, we performed an odd/even split of experimental scanning blocks to cross validate all core findings.

Insight into the nature of this functional dissociation within MPC can be found by considering parallel observations regarding network-based brain organization. While early functional connectivity research labelled the entire MPC as part of the ‘default mode network’^22^, subsequent work has instead separated recognition responsive regions of the MPC as falling within a distinct cognitive control^23,24^ or parietal memory^25^ network. This fractionation is consistent with the heterogenous anatomy of MPC^26,27^, which is comprised of multiple subregions, specifically the precuneus (PrC), posterior cingulate cortex (PCC), and retrosplenial cortex (RSC), that have variably been implicated in executive and episodic cognitive functions^17^ (Figure 1b). Together, a growing literature suggests that the observed dissociation of MPC responses during different types of episodic memory tasks may be better understood by considering the distinct types of mnemonic demands they require. For example, retrieval and representation of memory content predominates during autobiographical recollection, while control processes necessary for memory-based decisions are essential for recognition memory^28–30^. However, the considerable individual variation in functional and anatomical organization found in associative cortical regions like MPC makes standard neuroimaging methods that rely on spatial group-averaging suboptimal for detailed characterization of this region’s functional neuroanatomy^31^.

Advances in individualized, high-resolution neuroimaging have proven successful in better understanding the functional organization of other memory related structures, such the MTL^32,33^. Indeed, progress in applying these techniques to the MPC suggests great utility in capturing important features of functional organization which vary between individuals^34^. Specifically, recent developments in ‘precision-neuroimaging’ methods, where a small number of subjects are densely sampled through repeated scanning, not only improve the specificity of functional mapping within individuals, but also provide increased statistical power for conducting previously unfeasible analyses^35–38^. However, to date this approach has primarily focused on precision studies of resting state activity, but not functional task responses in MPC, particularly during episodic retrieval. As such, despite converging evidence linking the MPC with dissociable memory-supporting processes, precise mapping of these mnemonic contributions of the region is still lacking.

To this end, we analyzed an openly available functional magnetic resonance (fMRI) dataset^1^ collected at high-resolution using an ultra-high-field scanner (7T) while eight subjects performed thousands of stimulus-recognition decisions about semantically rich, naturalistic images over the course of many experimental sessions. This massive dataset afforded the statistical power and anatomic precision for evaluating MPC responses during item-recognition task performance that either reflected involvement in memory-based decision-making or were more indicative of mnemonic content representation. We operationalized this distinction by identifying MPC subregions where activity was associated with correct recognition-memory decisions or reflected the semantic content of stimuli. In doing so, we observed interleaving, though distinct, functional subregions of MPC where responses were sensitive to either recognition decisions or representation of the semantic content of stimuli, but rarely both. This striking functional dissociation was recapitulated in functional connectivity differences with the hippocampus, further supporting that MPC subregions act as members of distinct functional networks during mnemonic processing. Finally, we identified areas of MPC demonstrating categorically selective responses to person or place stimuli during recognition task performance, which were predominantly localized to semantically responsive regions. Together, our findings suggest a striking dissociation of mnemonic function exists within human MPC, which helps to better account for how distinct patterns of response may occur as a result of task demands during episodic retrieval and may reflect a common principle of organization throughout hippocampal-neocortical memory systems.

## Results

To examine the functional contributions of human MPC to recognition memory, we leveraged the Natural Scenes Dataset (NSD^1^), which comprises high resolution fMRI data collected during an extended recognition memory paradigm as well as resting state (see Materials and Methods). In the recognition memory task, subjects were serially presented with images of natural scenes (photographs) and asked to perform a simple recognition decision of ‘old’ (seen image before) or ‘new’ (never seen image before) (Figure 1c). Importantly, this paradigm was performed across multiple scanning sessions, from which the first 12 sessions (totaling 9,000 trials per subject) were used for all reported task analyses (Figure 1d; see Materials and Methods). Together, the NSD provides a unique opportunity to examine the functional organization of human MPC during recognition memory decisions, with both high spatial resolution and robust statistical power.

### Functional neuroanatomy of item recognition and semantic processing in the MPC

Across sessions, subjects performed well on the recognition memory task (Figure 1d; mean accuracy = 80.34%, 95% CI = [76.55%, 84.21%]). We examined blood oxygen level-dependent (BOLD) fMRI responses within the MPC while subjects performed the stimulus recognition task to identify regions involved in successful recognition decisions and/or the representation of stimulus content during these decisions. MPC regions responsive to recognition decisions were identified as those where univariate BOLD responses were greater for correct recognition of old stimuli (hits) than for correct identification of new stimuli (correct rejections, Figure 1c,d,f). Whereas semantic regions within MPC were identified as those where BOLD activity patterns during recognition were correlated with a semantic model applied to each image (Figure 1e-f; see Materials and Methods).

Analyses revealed distinct recognition and semantic responsive regions within MPC for each subject. Strikingly, for all subjects, we found that these functional regions demonstrated minimal spatial overlap. However, the exact spatial configuration and extent of recognition and semantic regions within MPC varied across individuals (Figure 2a). For example, while recognition responses (hits > CR) for most subjects were observed in the dorsal PCC/RSC (dPCC/dRSC) and ventral precuneus (vPrC), the spatial extent of these clusters differed across individuals. In all subjects, recognition clusters were identified in the dRSC, while in 5 of the 8 subjects, reliable recognition responses were also identified in dPCC. As can been seen in Figure 2a, whether this dorsal recognition cluster was primarily located in the dPCC or dRSC varied from subject to subject, highlighting these subtle, but important, individual-differences in activation patterns. Additionally, in all but two of the subjects, a reliable recognition cluster was found in the middle of the PCC, around the splenial sulcus^39,40^. However, the exact location of this middle PCC recognition cluster was highly variable across individuals. Interestingly, when collapsing across individuals to perform a standard group analysis, only the dRSC and vPrC clusters survived thresholding (Figure 2b). These two group-level recognition clusters directly overlap with regions associated with the cognitive-control and parietal memory networks (CCN; PMN) within the MPC as identified via previous resting-state functional connectivity analyses^23–25,41^. Overall, these group data replicate prior observations of consistent MPC subregional engagement during successful recognition decisions, specifically coactivation of distinct dRSC/PCC and vPrC clusters. However, our ability to robustly quantify individual subject responses suggests such group maps greatly underestimate the degree of individual variability in MPC recognition responses.

**Figure 2:**
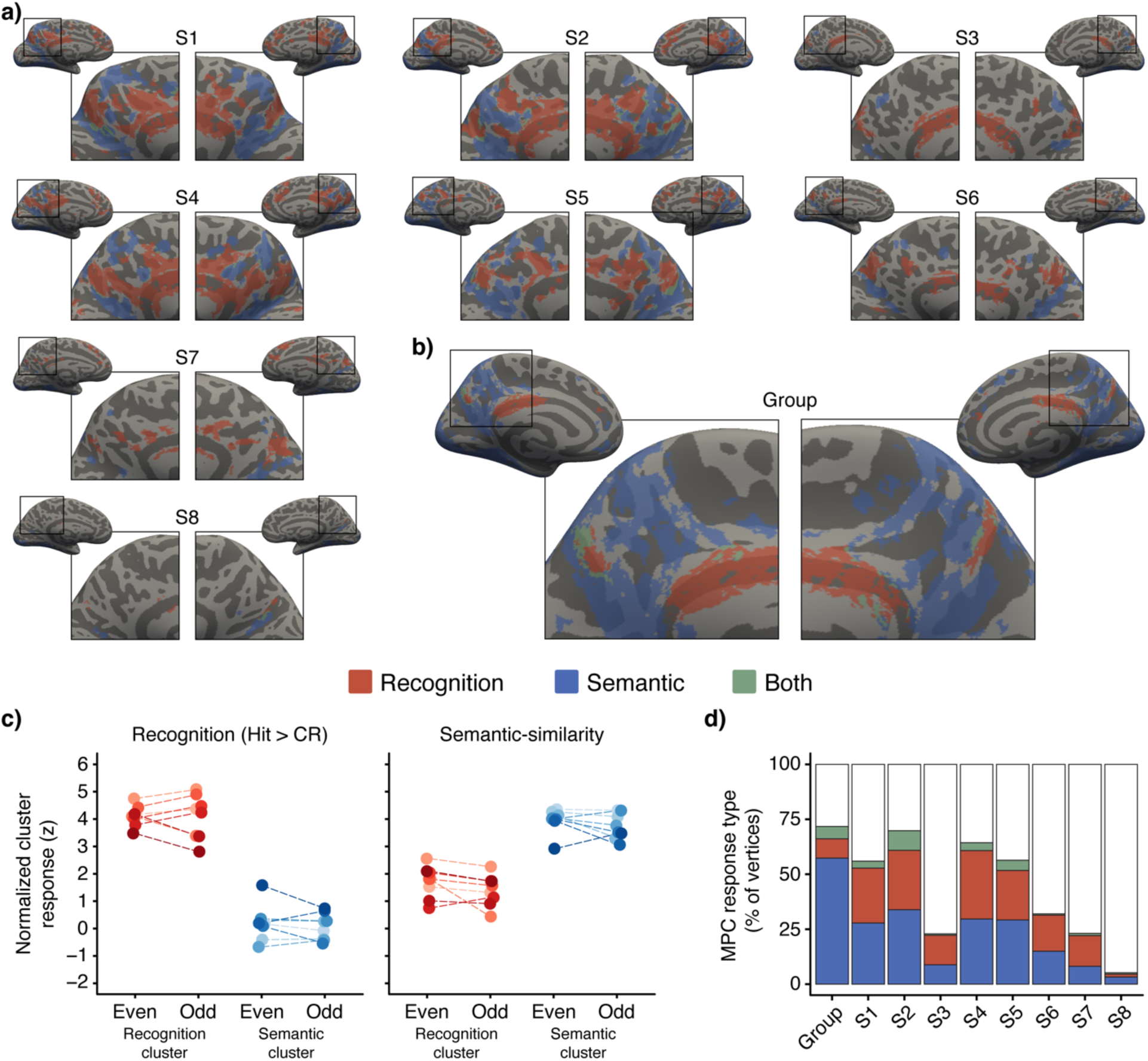
Stimulus recognition and semantic based responses within the MPC. **a**) Regions associated with recognition-decisions (red), image semantic content (blue), or both (green) visualized on the inflated medial surface anatomy for each individual. Depicted clusters were thresholded at z>3.3 (p<.0005) for all subjects except for S8 (z>3.1). **b)** Group map for the same contrasts and threshold as (a). **c)** Average recognition (left) and semantic (right) responses measured from recognition-clusters (red) and semantic-clusters (blue) from each subject. Scores were consistent across cross validation data-fold (even/odd). Left: Normalized BOLD responses to recognition decisions were significantly greater in recognition-clusters than in semantic-clusters. Right: Semantic responses were significantly greater in semantic-clusters than in recognition-clusters. Colors and dashed lines represent individual subjects. **d)** Mean percentage of cluster type (recognition, semantic, or both) assigned to MPC surface vertices (both hemispheres) at the group level and for each individual.

Within MPC, responses associated with stimulus semantic content also demonstrated variability in spatial configuration and extent across individuals (Figure 2a). Importantly, unlike the univariate contrast used to identify recognition regions, semantic regions were identified based on displaying activity patterns during recognition that were correlated with a semantic model applied to each image (see Materials and Methods). In all subjects, responses associated with stimulus semantic content were observed within the parietal occipital sulcus. For 4 of the 8 subjects, this cluster also extended into the gyrus of the ventral PCC. For all but one subject, there was an additional semantic cluster in the dorsal precuneus (dPrC). Additionally, in some subjects a semantic cluster was identified around the splenial sulcus, anterior to the splenial recognition region noted above. Furthermore, in 5 of the 8 subjects, a semantic cluster was observed around the marginal ramus of the cingulate sulcus. Interestingly, semantic group maps showed widespread organization scattered through much of the MPC (Figure 2b). This group result indicates that while within-individual evidence for semantic content was below our clustering threshold, across subjects this subtle relationship was consistently present. Group maps of semantic responses overlap with those commonly observed for the canonical default mode network (DMN) via resting-state and task data^23,41^. However, depending on the parcellation scheme used, these semantic ROIs may also be considered to overlap with the contextual association network^42,43^ or distinct subnetworks of the DMN (e.g. DMN-A vs. DMN-B^31,44,45^). Additionally, semantic clusters generated here are similar to previous results from studies investigating not only semantic content^46^, but importantly autobiographical retrieval^47–50^. As such, our semantic group-level results reflect a network of MPC regions that have been previously identified during recollection tasks involving mnemonic representation.

Given the striking variability of MPC response patterns between individuals we sought to leverage the large number of trials in the dataset to ensure robust and reliable findings. To do so, we used a cross-validation approach, where task trial data was split in half (even/odd) and the analyses performed within each data partition. Consistency across the two data partitions was evaluated by mixed effects linear regression, revealing there was no main effect of data partition (odd/even runs) on analysis results (beta = 0.178, t(7) = 1.817, 95% CI = [-0.235,0.592], p = 0.112). Post-hoc tests revealed this lack of difference between data partitions was separately true for the recognition (t(7) = 0.680, mean = 0.086, 95% CI = [-0.213, 0.385], p = 0.519) and semantic analyses (t(7) = 1.987, mean = 0.271, 95% CI = [-0.052, 0.593], p = 0.087). The consistency of responses across separate runs of the task suggest that observed patterns of recognition-decision responses and semantic-content similarity scores were reliable and well-founded within individuals. Having established analysis results were consistent across runs (visualized in Figure 2c), the following analyses collapsed across partitions when appropriate.

Given the striking interdigitation of recognition and semantic clusters within MPC, we next sought to quantify the degree of spatial dissociation and functional specificity of these clusters. To do so, we compared the mean normalized responses of recognition and semantic clusters, within subjects, for both ‘recognition decision’ and ‘semantic similarity’ analyses. Specifically, we quantified the degree to which recognition clusters displayed recognition or semantic activity in the held-out data-fold and the degree to which semantic clusters displayed recognition or semantic activity in the held-out data-fold. We then collapsed these scores across the two data-folds. As expected, recognition responses were significantly greater in regions masked by recognition clusters than by semantic clusters (t(7) = 10.449, mean = 3.962, 95% CI = [3.066, 4.859], p < 0.001; Figure 2c). The opposite relationship was observed for responses extracted via the semantic analysis, whereby values masked within semantic clusters were significantly higher than those masked within recognition clusters (t(7) = –11.83, mean = –2.299, 95% CI = [-2.758, –1.839], p < 0.001; Figure 2c). As follows, there was a significant interaction between cluster-type and analysis type (t(28) = 17.460, beta = 6.261, 95% CI = [5.527, 6.996], p < 0.001; Figure 2c), indicating that response values were greater when the cluster-type matched the score-generating type. There was no main effect of response magnitudes being greater in the recognition analysis versus the semantic analysis (t(29) = 0.952, beta = 0.578, 95% CI = [-0.664, 1.820], p=0.349) nor was there any main effect of the magnitude of responses being overall greater across both analyses in recognition versus semantic clusters (t(29) = –1.370, beta = –0.8312, 95% CI = [-2.074,0.410], p=0.181). In summary, the combination of cross-run reliability and across-analysis dissociation demonstrates that subregions of the MPC were strongly biased towards either recognition or semantic responses.

Finally, to confirm the spatial dissociation between recognition and semantic regions in the MPC, we compared the percentage of surface vertices that were uniquely found within recognition, semantic, or both cluster types. As can be seen in the individual and group surface maps (Figure 2a, b), there was a striking lack of overlap between the two sets of clusters (Figure 2d). On average, 19% of MPC vertices were observed to have above threshold recognition responses, while 19% reliably demonstrated semantic content, with only 3% of vertices being implicated in both. Therefore, on average 59% of MPC vertices demonstrated no reliable activity for either response type. Indeed, even at the group level, where a greater number of vertices were associated with semantic content than for any single individual (57%), the overlap between recognition and semantic vertices was still minimal (6%). These results further demonstrate the striking dissociation of interleaving recognition and semantic regions within MPC.

### Hippocampal subregion connectivity bias within functional regions of the MPC

Results from our previous analyses suggested a robust functional distinction between MPC regions that are responsive to specific recognition decisions from those that are responsive to the semantic content of recognition stimuli. This dichotomy can be framed as reflecting specific fine-grained details (recognition) compared with more broad conceptual information (semantics) relevant to memory behavior. Strikingly, a similar spectrum of functional specialization has been proposed to exist along the long-axis of the hippocampus. Specifically, the posterior hippocampus is thought to support fine-grained mnemonic details while the anterior hippocampus is proposed to support a more broad ‘gist’ representation^51^. Consistent with this, previous work using functional connectivity measures of resting state data have suggested vPCC regions that are part of the DMN (also termed contextual association network) are coupled with the anterior hippocampus, while dPCC regions that are part of the CCN/PMN are more strongly coupled with the posterior hippocampus^33^. Given these observations, we predicted a similar profile of hippocampal connectivity bias would be observed for our MPC recognition (greater posterior hippocampal coupling) and semantic (greater anterior hippocampal coupling) clusters.

To compare the hippocampal connectivity profiles of functionally defined MPC regions we independently calculated the average correlation of MPC recognition and semantic clusters to the anterior and posterior aspects of the hippocampus during both task and rest (see Materials and Methods). We subtracted the difference between hippocampal correlations for each cluster (anterior-posterior) to obtain a connectivity bias score, where negative values indicated stronger correlations to the posterior hippocampus and positive values indicated stronger correlations to the anterior hippocampus. Using mixed effects linear modeling, we first evaluated whether there was any effect of data-fold (even/odd), hippocampal hemisphere (left/right), MPC hemisphere (left/right), or any interactions therein, on the connectivity bias measure during recognition task performance. As this analysis showed no significant effects of these variables on hippocampal connectivity bias (all p > 0.5), we collapsed across data-fold and hemisphere to investigate the relationship between hippocampal bias and MPC clusters (cluster type: recognition/semantic) during task performance. Interestingly, there was a significant main effect of cluster type (t(7) = 6.289, beta = 3.577, 95% CI = [2.232, 4.922], p < 0.001), with recognition clusters consistently demonstrating a stronger connectivity bias towards the posterior hippocampus than semantic clusters. Post-hoc testing revealed that recognition cluster connectivity was significantly biased towards the posterior hippocampus (t(7) = –19.968, mean= –4.374, 95% CI = [-4.892, –3.856], p

< 0.001), while semantic clusters demonstrated a dissociable pattern of connectivity that was characterized by equivalent correlation strength across the length of the hippocampus (t(7) = – 1.301, mean = –0.797, 95% CI =[–2.246, 0.652], p = 0.235).

Next, we performed the same hippocampal bias analysis using resting state data (see Materials and Methods), with individual task identified MPC clusters serving as regions of interest (ROI). Recognition and semantic ROIs demonstrated statistically similar MPC-hippocampal connectivity whether they were generated from the even or odd data-folds (t(7) = 0.814, mean = 0.027, 95% CI = [-0.0513, 0.105], p = 0.442). Similar to task data, we observed a main effect of ROI type (t(7) = 9.590, beta = 2.221, 95% CI = [1.673, 2.768], p < 0.001), with recognition ROIs consistently demonstrating a stronger posterior hippocampal bias than semantic ROIs. Post-hoc testing revealed that for resting state data recognition ROIs demonstrated a significant connectivity bias to the posterior hippocampus (t(7) = –7.820, mean = –2.329, 95% CI = [-3.033, –1.624], p < 0.001), while semantic clusters demonstrated connectivity biased to a broader length of the hippocampus(t(7) = –0.745, mean = –0.108, 95% CI = [-0.451, 0.235], p = 0.481). Together, these connectivity results further support a striking dissociation between recognition and semantic regions of the MPC, one which extends into distinct profiles of connectivity with the hippocampus.

### Categorical selectivity to people and places in the MPC

In identifying a putative functional distinction between MPC regions supporting stimulus recognition decisions from those supporting stimulus semantic content, it is worth considering the features of this semantic representation, particularly as they relate to common elements of episodic memories. Recently, it has been observed that the MPC demonstrates categorical selectivity for people and place stimuli, common features of episodic memory, during perception and memory retrieval^52–57^. This observation is surprising, as historically the MPC has not been implicated in specific sensory processes, consistent with its general lack of direct connectivity with primary sensory regions^15^. However, if the semantic MPC regions identified in the analyses above are indeed involved in the broad representation of recognition stimulus content, then putative person and place selective regions should coincide within MPC semantic clusters. In addition, our findings would suggest that such an overlap is sensitive to performing mnemonic tasks.

The NSD provides a unique opportunity to examine both of these predictions, as it also includes a standardized visual category localizer task (fLoc) for each subject (see Materials and Methods). This dataset serves as a control for the main recognition task, by including unfamiliar, category specific cropped grayscale images (i.e. not natural images) presented during a non-recognition task (fixation with oddball ‘background only’ stimulus detection). It secondarily serves as a benchmark for identifying visual category selectivity within subjects. Therefore, to examine these questions we: i) mapped people and place selective regions across cortex in each subject using the standardized visual category localizer task (fLoc); ii) compared these functional maps with those constructed using only people and place stimuli from the NSD recognition task; iii) specifically compared the degree of overlap within the MPC for people/place maps generated via the fLoc and NSD tasks, with identified semantic clusters.

First, we mapped people and place selective regions across cortex in each subject using the fLoc task, where subjects were presented 10 visual categories, for which two categories were combined to define person (faces: adult and children) and place (houses and corridors) conditions (Figure 4a,c). Consistent with a large literature, person selective clusters were robustly and reliably observed in the lateral occipital, fusiform gyrus, and anterior temporal lobe (aTL) face areas^58–60^. Also consistent with prior work, place selective clusters were observed in the lateral occipital, parahippocampal gyrus, and parietal-occipital place areas^54,61^. However, no such robust evidence for person or place selective responses was observed in the MPC, where only sporadic, limited person and place selectively responsive clusters were observed outside of the parieto-occipital place area (i.e. medial place area). Using this established visual category localizer task, we were able to reliably map expected person or place selective cortices, however, we observed little evidence for person or place selectivity within MPC during perception.

**Figure 3:**
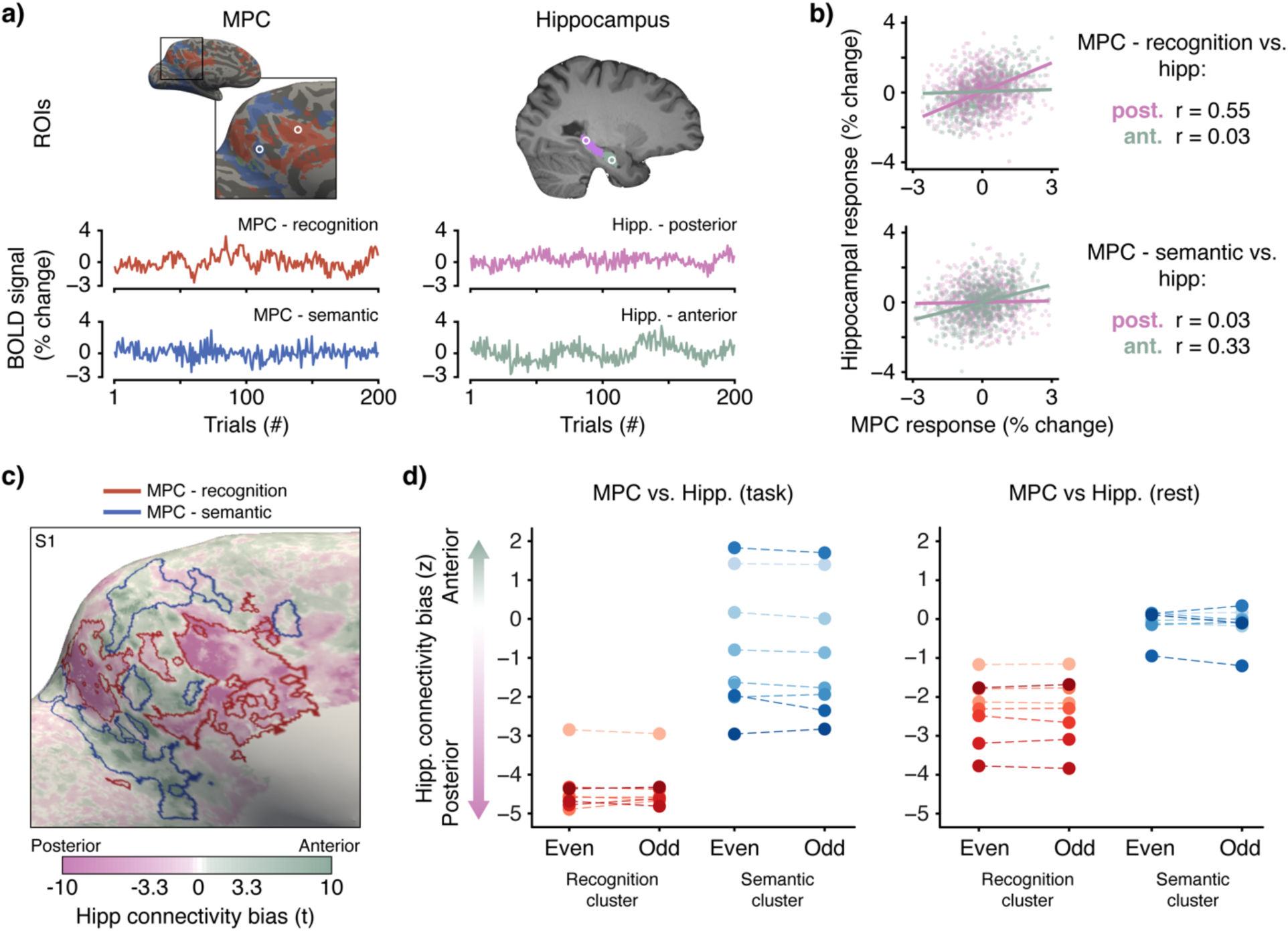
Functional connectivity between the MPC and hippocampal subregions during task and rest. **a**) Example beta-series of single voxels selected from a recognition cluster (red) and semantic cluster (blue) within MPC (left), as well as anterior (green) and posterior (purple) hippocampus (right) during one task session. **b)** Scatter plots show the correlation of beta-values (one session) from either the recognition cluster voxel (top) or semantic voxel (bottom), with both the anterior and posterior hippocampal voxels. **c)** Cortical surface map showing the across-session MPC-hippocampus connectivity bias values (t-scores) for S1. The surface color map indicates greater connectivity from MPC to the anterior (green) or posterior (pink) hippocampus. Outlines indicate the subject’s recognition (red) and semantic (blue) clusters used as ROIs. **d)** The average normalized (z-score) connectivity bias for voxels in recognition (red) and semantic (blue) ROIs from task (left) and rest (right). Voxels within recognition clusters demonstrated a significant connectivity bias to the posterior hippocampus, while no such relationship was observed for semantic cluster voxels.

**Figure 4:**
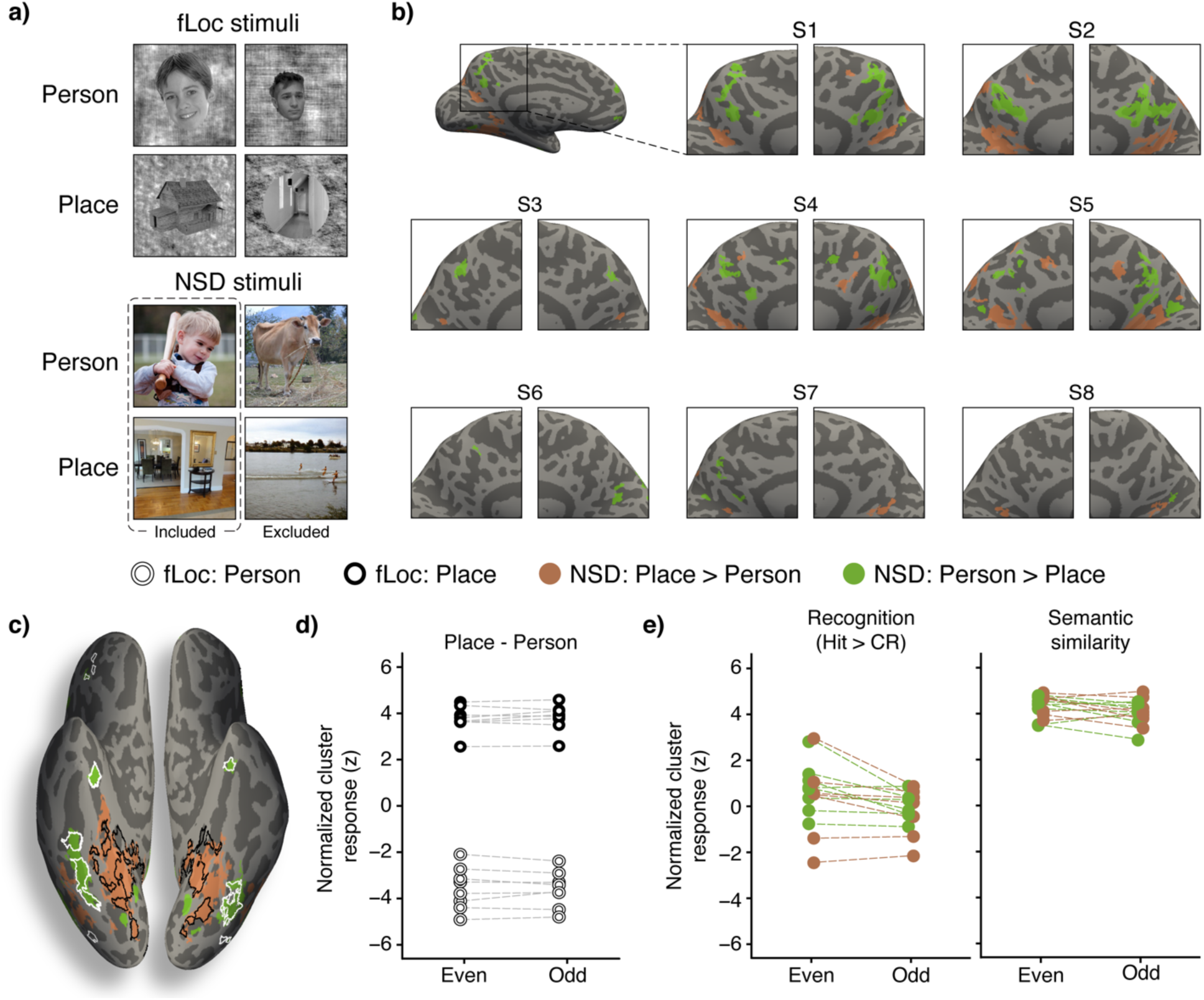
Categorical selectivity within the MPC. **a**) Top: Stimuli from the category functional localizer (fLoc) task from either the person (adult and child faces) or place (houses or corridors) categories. Bottom: examples of stimuli that were included (left column) or excluded (right column) as person or place stimuli in the NSD task categorical selectivity analysis. **b)** Regions demonstrating selective responses during the NSD task towards people versus places visualized on the medial inflated surface anatomy for each subject. **c)** Ventral surface visualization from one subject showing the spatial overlap of person (white outline) and place (black outline) clusters from the category localizer and person (green) and place (orange) clusters from the NSD task. **d)** Person (white) and place (black) selective clusters from the category localizer demonstrated similar selectivity to person (negative values) and place (positive values) during the NSD task. The categorically biased responses were similar across even and odd runs of the NSD-task (x-axis). **e)** Person and place clusters from the NSD task were used as masks for the recognition and semantic analysis. While there was no significant recognition response found within person or place clusters overall, significant semantic content responses were observed in both cluster types. Throughout, green is used to indicate person clusters and orange indicates place clusters identified from the NSD recognition task. NSD-task clusters were thresholded at z>3.3 with at least 20 contiguous vertices. Category localizer clusters were thresholded at t>3.3 and at least 20 contiguous vertices.

Next, we performed a similar analysis using selected person and place stimuli from the NSD recognition task, to benchmark the identification of person/place selectivity against maps generated from the fLoc task above (see Materials and Methods). With this goal in mind, we compared responses to person/place stimuli during the recognition task within person/place clusters identified via the fLoc task. As expected, person and place clusters generated from the category localizer (fLoc) analysis also showed categorically selective responses to images featuring people or places during the NSD task. Specifically, we found that responses in category localizer person and place clusters were significantly different for person versus place stimuli during the NSD task (paired t-test: t(15): 24.208, mean = 7.361, 95% CI = [6.713, 8.009], p < 0.001), with person clusters generated from the fLoc task demonstrating high person selectivity during the NSD task (t(7) = 18.531, mean = 3.786, 95% CI = [3.302, 4.269], p < 0.001), and responses within place clusters generated from the fLoc task being significantly selective in the opposite direction towards place stimuli (t(7) = –11.900, mean = –3.575, 95% CI = [-4.286, –2.865], p < 0.001). These results confirmed a strong consistency in cortical regions identified as person or place selective via a well-controlled localizer task or person/place trials of the NSD recognition task. However, while there was strong overlap in the maps of person/place selectivity across both tasks, only the recognition task included robust clusters within MPC. These observations support the efficacy of identifying person/place selective regions via the NSD recognition task and also suggest that selectivity to these categories within the MPC may be sensitive to mnemonic judgements.

In light of this task difference, we next sought to map the functional organization of people/place selective regions within MPC across individuals. While the location of selective regions varied greatly between subjects, certain patterns did emerge (contrast maps are shown in Figure 4b). For example, while all eight subjects exhibited at least one person-selective patch within the MPC in both hemispheres, the location, number, and extent of person selective regions varied from subject to subject. Across individuals, person selective clusters differed in whether they were found in the parieto-occipital sulcus, splenial sulcus, or near the marginal ramus. Regarding places, all but one subject exhibited a selective region in the inferior parieto-occipital sulcus. For most subjects, this cluster did not extend into the neighboring gyrus of the vPCC. Together, we observed distinct person and place selective regions within the MPC, consistent with prior observations^53^, however, our precision data suggested these maps showed anatomical variability across subjects, much like identified semantic clusters.

To quantify functional overlap between the person and place selective regions of the MPC with the recognition and semantic clusters identified above, we conducted a mixed effects linear regression, comparing recognition and semantic content responses from within identified person and place clusters of the MPC. Across both person and place clusters, we found a main effect such that average responses were significantly higher for the semantic analysis than the recognition contrast (t(47) = 17.424, beta = 4.049, 95% CI = [3.573, 4.524], p < 0.001). Post-hoc tests revealed that overall the average recognition responses within person and place clusters did not significantly differ from zero (t(7) = 0.756, mean = 0.252, 95% CI = [-0.537, 1.042], p=0.474; Figure 4e), nor did recognition responses significantly differ between person and place clusters (t(13) = 1.138, mean = 0.341, 95% CI = [-0.307, 0.990], p = 0.276). Conversely, semantic responses from the same person and place clusters were significantly greater than zero (t(7) = 32.12, mean = 4.188, 95% CI = [3.880, 4.497], p < 0.001), and were similar between person and place clusters (t(13) = –0.770, mean = –0.123, 95% CI = [-0.468, 0.222], p = 0.455).

Finally, we looked at the degree of overlap between categorically selective clusters in the MPC (collapsing across both person and place selective clusters) with recognition and semantic clusters (recognition task data). After averaging across data-fold and hemisphere within subjects who had at least one person or place selective MPC cluster, we found that 75% of the person-selective cluster vertices and 69% of the place-selective cluster vertices overlapped with semantic clusters, while only 1% of person-selective and 3% of place-selective cluster vertices overlapped with recognition clusters. A remaining 19% of person-selective and 24% of place-selective cluster voxels fell outside of either the recognition or semantic clusters, while the remaining 5% of person– and 4% of place-selective vertices overlapped with clusters that demonstrated both recognition and semantic responsivity. Together, these data further support the view that identified semantic clusters within MPC contain subregions displaying selectivity to person or place stimuli, specifically when making mnemonic based judgements.

## Discussion

We analyzed a high-resolution fMRI dataset collected while subjects performed thousands of stimulus-recognition trials to characterize the functional contributions of MPC to recognition memory. Within individuals, we were able to identify two interleaving, though largely non-overlapping, mnemonic subnetworks. One subnetwork demonstrated responses indicative of supporting recognition memory decisions, while the second reflected the semantic content of mnemonic stimuli. These two subnetworks were further differentiated by their profile of functional connectivity with the hippocampus and responsivity to person and place stimuli. Recognition regions demonstrated biased connectivity with the posterior hippocampus and no categorical selectivity to people or places. Conversely, semantic regions were more strongly correlated with the anterior hippocampus than recognition regions and did demonstrate categorically selective responses to people and places. Importantly, we observed robust idiosyncrasies in the spatial organization of recognition and semantic regions within individuals that differed substantially from group maps. Together, these results revealed a striking dissociation of mnemonic function within human MPC which, as detailed below, converges with multiple lines of evidence highlighting similar dichotomies of functional organization throughout putative brain networks supporting episodic memory.

Within individuals, we observed separate, though closely interdigitated, recognition and semantic responsive MPC regions. Recognition responses were consistently found in the dRSC and dPCC, as well as the vPrC and near the splenial sulcus. Meanwhile, semantic regions were observed within the parieto-occipital sulcus, vPCC, mid PCC, and dPrC. At the group-level, recognition regions were limited to the dRSC and vPrC, mirroring previous results from item-recognition studies^25,62^. Group-level semantic clusters were observed from the parieto-occipital sulcus and vPCC to the dPrC, reflecting similar results that have been found in analyses of semantic representation^63–65^ and episodic recollection^13,47,66^. Additionally, this group-level functional neuroanatomy mimics dissociations reported in studies directly comparing MPC responses to item-recognition against autobiographical retrieval^20,21^, as well as studies comparing content learned in a laboratory setting versus lived experience^67,68^. Furthermore, similar dissociations have been observed in studies examining episodic recollection versus familiarity^69,70^, which have led to the proposal that patterns of activity in the MPC are related to the level of cognitive control versus mnemonic representation involved during retrieval^66^. Indeed, converging evidence from human and animal studies suggests that MPC recognition regions may serve a broader executive function, not limited to mnemonic processing (for review see^17^). For example, activity in the dPCC/dRSC and vPrC have consistently been linked with value and decision-outcome processing^71–75^, environment updating processes^76,77^, event segmentation^78^, decision-risk evaluation^79^, goal-congruency^80^, explore vs. exploit decisions^81^, saliency processing^82^, and executive control^28,30^. These dPCC/dRSC results associating the subregion with executive or decision-making processes are in stark contrast to neuroimaging studies implicating the vPCC, which largely link this subregion to the recollection of experiences^13,83,84^ or scene construction^85,86^. In addition to task-based findings, previous analyses of resting-state BOLD activity have been used to identify dissociations in functional network involvement between MPC subregions that closely align with the recognition and semantic clusters we observed^23,41,42,87^. Recognition clusters overlap with the frontoparietal or parietal-memory networks, associated with cognitive control and familiarity processing^25,88^, while semantic clusters overlap with the default mode network, classically associated with autobiographical memory and internally directed cognition^7,89^. Our results converge with growing evidence that a functional dissociation exists within the MPC during episodic recollection and recognition, which distinguishes regions involved in executive control during memory-based decisions from regions involved in the retrieval and representation of mnemonic content. Importantly, while only examining recognition behavior, our identification of both recognition and semantic subregions during this task highlights how retrieval task demands will differentially drive distinct univariate patterns of MPC response, which may fail to fully capture other, more subtle, task relevant activities throughout the MPC.

What factors may help account for this dissociation of functional regions observed in MPC? Interestingly, we found that recognition and semantic clusters demonstrated distinct patterns of functional connectivity with the hippocampus during task performance and rest. Specifically, we observed that recognition clusters exhibited a strong connectivity bias towards the posterior versus anterior hippocampus. Conversely, semantic clusters were observed to have a broader connectivity bias between the anterior and posterior aspects of the hippocampus, and consistently weaker posterior connectivity strength than recognition clusters. Given that such task/rest coupling often conveys putative functional association^90–93^, recent progress in revealing a hierarchy of organization throughout the longitudinal axis of the hippocampus can serve to help better understand the nature of our dissociations within MPC. For example, during memory retrieval, the anterior hippocampus is associated with broad, gist processing involved in autobiographical retrieval, while the posterior hippocampus is linked to more specific, detailed representations in support of goal-directed processing^33,51,94^. More generally, the hippocampus is thought to be characterized by a gradient of representational granularity that shifts from broad to more fine-tuned representation along the anterior to posterior axis, respectively^95–97^. An interesting parallel can be found in another key cortical region for episodic memory, the lateral parietal cortex. There, dorsal regions, such as the intra-parietal sulcus, show stronger connectivity to the posterior hippocampus and parietal memory/cognitive control networks, and are implicated in item-recognition and familiarity responses. Conversely, the inferior parietal lobule, specifically the angular gyrus, displays biased connectivity to the default mode network, and is implicated in autobiographical and item recollection^20,66,98–101^. These converging results across multiple regions suggest an important dimension of executive versus mnemonic processing that shapes the functional organization of brain structures supporting episodic retrieval.

The recognition and semantic subnetworks of the MPC are further differentiated by their selective responses to categorical stimuli. Using the naturalistic stimuli from the NSD, we found categorically-selective responses in the MPC when individuals were making memory decisions for complex, naturalistic stimuli. Though MPC has not traditionally been included in descriptions of specific sensory processes, a growing body of literature has linked MPC to spatial processing and egocentric navigation^102–104^, including categorically selective responses to person and place stimuli^55,61,83,105^. Our findings build on previous reports of categorically selective responses reported in MPC during recognition of experimenter-controlled people and place stimuli and during recall of highly familiarized memories of famous or personally relevant people and places^53,83,106^. Indeed, previous work has found that categorically selective responses in the MPC during retrieval are modulated by the degree of familiarity one has with a given stimulus. For example, studies have found greater responses to personally familiar versus famous people^53,56^, and also greater responses to famous and familiar people than unfamiliar people^52,107^. We found that, outside of the parieto-occipital sulcus, categorically sensitive responses were not observed during a passive localizer in the MPC, adding to the evidence that the responses in the region are mnemonically sensitive. We also confirmed that categorically selective responses were predominantly located within semantic and not recognition clusters, serving to further support the role of semantically responsive MPC regions in broad, categorical representation, dissociated from the more executively implicated recognition regions. Together, these findings integrate recent observations of categorically specific representation within MPC during retrieval, suggesting that identified person/place regions are part of a larger semantic region representing retrieval contents. However, these data also suggest that features of particular relevance to episodic memory, like the people and places that make up our social experiences, maybe more represented over other feature domains, reflecting a hippocampal transformation of ventral stream sensory inputs into the MPC. We leveraged the large amount of within subjects data of the NSD to perform previously unfeasible analyses in order to examine the complex organization of functional subregions within the MPC. However, there are some aspects of the experimental design that limit the scope of conclusions we can draw. For example, to allow for a large number of trials subjects were only required to make a binary old/new response to each serially presented stimulus. As each stimulus remained on the screen during the decision period, we cannot robustly dissociate perceptual versus mnemonic effects during recognition, and as such, it is possible that identified semantic clusters are more perceptual in nature. However, the lack of robust MPC responses to controlled people/place stimuli during the visual localizer task (which lacked recognition) compared to the clear responses observed during the recognition task, suggests that the semantic MPC activity observed here is more likely related to mnemonic than perceptual processes. While new experiments are beyond the scope of this work, neural networks trained on image categorization (e.g. AlexNet^108^) to extract low-level visual features^109–111^, may provide a fruitful avenue for dissociating perceptual and mnemonic/semantic responses with in MPC. It is also possible that further subdivisions exist within the recognition and semantic MPC regions that we cannot parse using the binary old/new decisions made by subjects. For example, subregions responsive to recognition decisions within lateral parietal cortex further show a graded response magnitude based on the confidence/strength of recognition judgements^98^. Similarly, subdivisions in the type of information represented may exist within semantic MPC regions. While were able to consistently identify general semantic regions using a complex natural language processing model (Google USE) and indeed identify person and place selective regions, richer evidence for functional specificity^112^ may be more easily observed in a task requiring recollection of dynamic episodes.

To build on our understanding of the role of MPC in episodic memory and cognition in general, future work should continue to precisely characterize the functional neuroanatomy of the region. For example, while the present work indicates one set of subregions supports executive processes during episodic memory tasks, further research is needed to determine whether this is the same region that is implicated in broader decision-making processes like subjective-value or decision-outcome evaluation^71,72,74^. Better identification of the neural substrates that support seemingly disparate cognitive functions will improve our understanding of the relationship between cognitive control and memory driven behaviors. Similarly, processes involved in different types of memory retrieval like evaluating implicit familiarity signals, conducting memory search, or visualizing episodes may involve a complex interaction of executive and mnemonic functions, differentially supported by distinct subregions of the MPC. To better elucidate the unique contributions of MPC, it will be beneficial to compare how memory content representations differ from other relevant brain structures^113^. Future work will also benefit from comparing across a larger battery of memory tasks to test how these retrieval process requirements modulate different forms of MPC engagement. Furthermore, our results suggest that the use of precision-neuroimaging approaches will be essential for providing the spatial resolution and statistical power necessary for accurately mapping these complex functional dissociations. In all, the results reported in this work suggest that consideration of individual differences in MPC functional organization will allow for better characterization of the region’s contributions to memory and more general cognition.

Results from the current study represent an important step towards more precisely understanding how the MPC contributes to episodic memory and cognition more generally. Broadly, we show that a stark dissociation exists between interleaving subregions of the MPC involved in processing of recognition-decisions versus semantic content at the individual level. Converging evidence suggests that recognition regions support fine-grained evaluation of stimuli in support of mnemonic decision-making. Conversely, semantic regions are involved in representation and retrieval of broad details during episodic retrieval. While some theories suggest a role for the MPC in executive and evaluative function^114^, other theories focus on a mnemonic role for the region^115^. Our results suggest that both sets of contributions are performed by the MPC, though interleaving sets of regions are primarily responsible for either function. Therefore, the commonly observed dissociation of episodic retrieval task activity within MPC is likely driven by the degree to which executive versus mnemonic components are required, and that rather than two separate sets of processes, these executive and mnemonic components operate in complex interaction within MPC as a convergent zone for such neural systems^17^. These functional attributes may help to better understand the unique contributions of the MPC, while more broadly revealing a principle of functional organization common throughout memory systems in the human brain.

## Materials and Methods

The data analyzed here were collected and initially preprocessed as part of the Natural Scenes Dataset (NSD^1^). The NSD is a large-scale fMRI dataset collected using ultra-high field fMRI (7T) while subjects completed hundreds of stimulus recognition decisions on semantically rich, naturalistic images (from the COCO dataset^116^). The NSD includes preprocessed, high-resolution functional images (beta-maps with upsampled 1mm-isotropic voxels and robust hemodynamic response function fitting), but also fMRI data from rest and visual localizer tasks. For a full description of the dataset, as well as specific methods involved in collecting and preprocessing the original data, please see the original manuscript^1^. Below we outline the analysis steps relevant to the current manuscript.

### Subjects

Eight subjects from the University of Minnesota community were included in the experiment (two male and six female, ages 19-32). All subjects had normal or corrected to normal vision, and no known cognitive deficits. Informed written consent was collected for all subjects, and the protocol was approved by the University of Minnesota institutional review board.

### MRI acquisition and preprocessing

Anatomical data was collected using a 3T Siemens Prisma scanner and 32-channel head coil. For the analyses included in this manuscript we used the preprocessed, 1mm preparations of the volumetric anatomical images and the corresponding Freesurfer output for surface spaces. Additionally, we used the provided transformation matrices and code to map between individual volumetric and surface spaces, as well as to normalize functional outputs from individual surface space to the fsaverage surface. Task and resting-state functional scans were collected using a 7T Siemens Magnetom passively shielded scanner using a 32-channel head coil. For functional analyses, we used the 1.0mm preparation of the functional timeseries data (described as beta-2 in the original manuscript). Briefly, this preparation is characterized by treating each trial as an independent ‘condition’ and fitting each voxel with a best-fitting hemodynamic response function (HRF) determined for each voxel and session. This is done by using a cross-validated approach for fitting a library of potential HRFs to each voxel, choosing the function with the best fit, and using that function to evaluate the entire session for that voxel. Next, these preprocessed HRF-fit volumetric timeseries were resampled via cubic interpolation onto the subject-native surface at three different depths (layers 1-3), and then averaged across all depths to create a single surface timeseries matrix (vertices x trials). These matrices were used for all subsequent functional analyses except for hippocampal timeseries data, which was taken from the corresponding volumetric timeseries (pre-surface interpolation voxel x trial matrices). No smoothing was applied to these timeseries. To return data from int16 minimal-storage format to percent signal change for analyses, all vertex values were divided by 300. Then, values were normalized by z-scoring within each vertex and session.

### Task details

Subjects performed a continuous image recognition task, where on each trial they were shown an individual image and asked to respond as to whether the image was identical to any image that had been presented previously during an experimental session (‘old image’) or had not been presented during any experiment session (‘new image’). The experiment was organized such that each scanning session contained 12 runs and 75 trials were presented per each run. The first three and last four trials of each run were blank, with five additional blank trials being sporadically presented within each run (time between blank trials 9-14 trials). Additionally, the 63^rd^ trial of even runs was blank, leading to a total of 750 task-trials per session [(63 trials *6 odd runs) + (62 trials *6 even runs) = 750]. Subjects completed between 30-40 scan sessions each, taking part in experimental sessions approximately once a week over the course of a year. Depending on the number of sessions completed over the course of the entire experiment, subjects performed between 22,500-30,000 trials. Each subject was shown up to 10,000 unique images, and each image was repeated up to three times. Across all subjects, 73,000 unique stimuli were employed, with 1000 stimuli chosen to overlap between all eight subjects.

The NSD stimulus recognition task was designed so that individuals continuously made memory decisions on hundreds of visual stimuli across a wide range of inter-repetition gaps. Each trial lasted for 4s, wherein a single stimulus and a small, semi-transparent fixation dot were presented for 3s, followed by a 1s inter-stimulus-interval during which only the fixation dot remained on the screen. Subjects were instructed to maintain central fixation and to respond by pressing 1 with their index finger for ‘new’ images or press 2 with their middle finger for ‘old’ images. Subjects were allowed to change their response as many times as they wanted during the trial window, and accuracy was defined by their last button press. ‘Old’ trials could be those where stimuli had been shown either previously during the current session (‘easy-old’ trials) or during a previous session (‘hard-old’ trials). The task was designed so that as subjects performed the experimental sessions over time, the number of old items increased, the ratio of easy to hard trials decreased, and the ratio of new to old trials decreased. As such, a cutoff of trials to be included in subsequent functional analyses was chosen such that the ratio of old to new trials, as well as the ratio of hard to easy trials, remained below 1.5:1. Importantly, the shift in ratio of trial types corresponds to a similar drop in recognition performance. We evaluated adjusted hit rates for easy and hard old trials (the rate of correct recognition decisions for easy or hard trials divided by the overall false alarm rate^1^) for sessions which all subjects performed and were publicly available (sessions 1-26). Performance was significantly worse for both easy and hard old trials after compared to before the 12^th^ experimental session (easy: t(199) = –10.71, b = –0.140, 95% CI = [–0.165, –0.113], p < 0.001; hard = t(191) = –10.995, b = –0.109, 95% CI = [-0.129, –0.090], p < 0.001). This cutoff, set at including sessions 1-12, considered both the trial-type ratio and subject performance, and was chosen in order to maximize generalizability of the results from the current analyses while still taking advantage of the incredible statistical power to evaluate memory decisions afforded by the NSD task design.

## Analysis Details

### Recognition-decision and semantic-content analysis overview

Our first set of analyses were aimed at identifying regions of the MPC where activity was associated with recognition memory decisions and/or semantic content. To ensure robust results, a cross-validation sampling method was used. The z-scored cortical surface data was first divided into even and odd runs from the first 12 fMRI sessions. Then recognition-decision and semantic content analyses were performed to separately create clusters on one set of the data (e.g. even trials), and to test cluster analysis scores on the held-out set (e.g. odd trials). This process was repeated once, so that each partition of data served once as the cluster-generating dataset and once as the score-generating dataset. All analyses were performed in Python 3.7 using custom code from NiLearn, Scikit-learn^117^, NiBabel^118^), and SciPy^119^ packages, as well as code provided by the NSD researcher team (https://github.com/cvnlab/nsdcode).

### Recognition decision analysis

In order to identify regions of the MPC where activity was related to recognition decisions, we first performed a univariate analysis of BOLD activity during hits versus correct rejections. At the first level of this analysis, the normalized beta values within each session for each subject were compared between hit recognition trials (correct ‘old’ responses) and correct-rejection trials (correct ‘new’ responses), generating a by-session t-score map. Next, we tested whether differences in activity between hits and correct rejections were consistently greater than would be expected by chance by normalizing by-session t-score maps (into z-scores), and running a one-sample t-test comparing the by-session z-scores against an empirical mean of 0 for each vertex. The corresponding t-scores from this second level analysis were normalized (to z-scores) for thresholding and comparison across subjects. In order to control for multiple comparisons without drastically decreasing analysis sensitivity, cluster thresholding was performed by including only clusters where at least 20 contiguous vertices were above a z>3.3 (p<0.0005) threshold. For one subject (S8), a threshold of z>3.3 led to no surviving clusters. For this subject, the threshold was lowered to z>3.1 (p<.001) so that we could run a more complete statistical analysis of reliability across individuals. These thresholded maps were used for visualization and characterization of recognition-responsive regions of within the MPC.

### Semantic content analysis

We also sought to identify regions of the MPC where BOLD activity was related to the semantic content of recognition-stimuli. In summary, semantic content regions were identified as those where activity patterns during recognition were correlated to a semantic model applied to each image (Google USE_v5^120^). This analysis involved a two-component surface searchlight approach. First, for each stimulus a semantic feature vector was generated from the annotation captions included for each image in the COCO dataset (more details on the caption collection and validation can be found in the original manuscript^121^). The annotations were five unique human-generated sentences describing each stimulus (e.g. ‘a boy in a grey sweater holding a wooden baseball bat’). These sentences were converted to quantitative semantic feature vectors for each trial, by concatenating all five sentences and feeding them into Google’s Universal Sentence Encoder. The Google USE_v5 encoding model was selected as it is optimized for multi-word semantic evaluation. The result of feeding annotations through the semantic model was a 512 semantic feature-vector for each stimulus. The dissimilarity between each stimulus’s semantic vector was measured using cosine distance. This generated a stimulus x stimulus sized matrix of semantic dissimilarity for each stimulus seen during the included trials (e.g. even/odd runs) of each session.

The second step of the semantic content analysis involved generating a corresponding dissimilarity matrix for stimuli based on BOLD patterns of activity. A surface searchlight approach was used to generate these neural dissimilarity matrices. For each vertex on the surface, a 3mm radius searchlight sphere was generated (mean vertices = 90) and converted into a vector. Cosine distance of BOLD activity in response to each stimulus was calculated to generate a vertex-centered dissimilarity matrix for each searchlight sphere across all included trials. Lastly, the relationship between the neural dissimilarity matrix and the semantic-model’s dissimilarity matrix was quantified by taking the Pearson correlation between the two matrices. The output correlation value was assigned to the sphere center as the semantic similarity score for that vertex. This process was repeated for all vertices, across all sessions, and within each subject in the same cross-validated manner as above. Having computed the by-vertex, within-session correlations between neural activity and the semantic model, we next sought to determine how reliable this relationship was across sessions. Quantification of semantic content reliability was done by normalizing correlation values (Fisher transforming) and then computing a single sample t-score to evaluate statistical difference from zero for correlations across sessions. Next, the resulting semantic reliability maps (t-scores) were normalized (to z-scores) and cluster thresholding of z>3.3 (equivalent to p<.0005) and cluster size minimum of 20 contiguous vertices was applied. In order to mirror the memory-decision analysis, this cluster threshold value was lowered to z>3.1 (p<.001) for S8 (the same subject as above).

We next investigated group-level results for both the recognition and semantic content analyses so that we could compare findings to previous research. Here, pre-thresholded second-level z-score maps from each analysis for each subject were normalized to the fsaverage surface using the ‘NSD_mapdata’ function and publicly available NSD transformation matrices. Group-map t-scores were computed within each vertex, and then converted to normalized z-scores for thresholding and visualization.

A cluster-based analysis was next carried out to quantify how functional responses differed between MPC clusters implicated in the recognition-decision and semantic content analyses within individuals. Using the cross-validation method outlined above, clusters were generated using either even or odd runs across the first 12 sessions. Then the held out set of runs was used to generate score maps for each subject (e.g. strength of recognition or semantic content responses). This scheme was repeated once so that each set of data was used once to generate clusters and once to generate z-score maps. To check whether analysis scores were consistent across the two cross-validated data-folds, we used mixed effects linear regression with the difference in cluster-scores between even and odd runs as the fixed effect and a random effect of subject intercept. Having established scores did not significantly differ across folds, we averaged results across folds, and then quantified the degree of overlap between regions implicated in the recognition versus semantic content analyses. To do so, we took the four output values for each subject, representing a factorial combination of cluster type (recognition or semantic content) by analysis-map (recognition or semantic content), and input them into a mixed effects linear regression model that also included a random subject intercept term and fixed effects of cluster type and analysis-type. Results from this analysis are visualized in Figure 2c.

We also evaluated the degree of overlap between recognition and semantic content regions by quantifying the percentage of MPC vertices that we observed to be above threshold for the recognition, semantic content, or both analyses. This value was determined by taking the average number of vertices that were above threshold and then dividing by the total number of MPC vertices (visualized in Figure 2d). These analyses allowed for us to gain insight into the degree and arrangement of processing overlap for recognition and semantic content within MPC subregions.

### Category localizer (fLoc) experiment analysis

The functional localizer, developed by the Grill-Spector lab (fLoc^122^; http://vpnl.stanford.edu/fLoc/), was used to examine neural responses to isolated visual object categories. This functional localizer has been previously used to identify regions involved in face or place processing^122^ and involved displaying curated images from 10 categories to subjects. The stimulus categories were either characters (words or numbers), bodies (bodies or limbs), faces (adult or child), places (houses or corridors), or objects (cars or instruments). The images used in these stimuli have been carefully selected and cropped to display only features in the category of interest. Stimuli were presented in grayscale on scrambled grayscale backgrounds, and adapted to fill an 8.4° x 8.4° square and with a semi-transparent fixation dot centrally located, mirroring the NSD stimulus presentations. Stimuli were displayed in 8 trial mini-blocks with each stimulus being presented for 0.5s each, totaling 4s per mini-block. Each run contained six presentations of each of the 10 categories, with each run lasting 300s total. Six runs were collected, totaling 36 mini-blocks and 288 image presentations per category.

The NSD includes t-score contrasts from the category localizer task, reflecting categorical selectivity across individual cortical surfaces. These selectivity maps were generated using GLMdenoise and a ‘condition-split’ strategy (for more details see original manuscript and supplementary files). For person selectivity, t-scores were generated by comparing betas for face stimuli (adult and child) to all other categories. Similarly, place selectivity was computed by comparing house and corridor responses to all other categories. We then generated selective clusters by taking these precomputed t-score maps, thresholded at t>3.3, and applied a cluster threshold of necessitating at least 20 continuous vertices. These clusters were then applied to the NSD-based categorical selectivity analysis described below.

### Categorical representation analysis

While much of previous research examining categorical selectivity has used experiment-controlled stimuli, like the functional localizer above, the NSD’s use of the COCO stimuli allows for examination of person and place selective processing in the MPC across a wide range of naturalistic images. Our first step in performing a categorical selectivity analysis, was to curate NSD stimuli to those that primarily featured either people or places. We used pre-generated bounding boxes (https://github.com/cocodataset/cocoapi and https://github.com/ACI-Institute/faces4coco) to establish the percentage of each image that was occupied by a person’s body or face, or by any objects or animals within the stimulus. If at least 10% of the image was a face or 15% of the image was a person’s body (face, body, or limbs), we considered that image to be highlighting an individual, and to be a ‘person’ image. Conversely, to select ‘place’ images, we performed the opposite inclusion steps. If the stimulus included at least 15% of any given object (animal, food, etc.…) or any amount of a person, we determined that the stimulus featured an object, person, or animal and as such removed that stimulus from the ‘place’ image set. All other stimuli were included. This stimulus selection process allowed for a robust examination of categorical selectivity within the MPC by including complex naturalistic stimuli that featured either people or places, but also contained other objects or actions. Examples of included and excluded stimuli can be seen in Figure 4a.

Next, we quantified categorical selectivity within the MPC by evaluating the same preprocessed data as used in the previous recognition and semantic content analyses (the beta version-2 timeseries surface data from sessions 1-12 of the NSD recognition task). In similar fashion to the previous recognition and semantic analyses, we conducted this analysis in a cross-validated manner. Whereby, one half of the data (e.g. even runs) was used for analyses and then the process repeated with the held out set of data. By-session bias scores for either person or place stimuli (t-scores) were generated by comparing betas from our curated person versus place stimuli. Next, the by-vertex statistics of categorical bias were quantified by conducting a single-sample t-test to compare the bias score across sessions against empirical chance of zero. Across session categorical selectivity bias maps (t-scores) were normalized to z-scores for thresholding and visualization. For cluster generation, cluster thresholding was performed to the second level categorical selectivity maps from the NSD task (z-scores) analysis to only included vertices where z>3.3 (p<.0005), and which were part of a group of at least 20 contiguous vertices. For visualizing patterns of categorical selectivity in the MPC within individuals (Figure 4b), selectivity maps from each data fold were averaged together, and thresholding applied to those averaged maps.

To confirm that this analysis was capturing similar aspects of stimulus processing as the well-controlled category localizer, we compared results from the localizer and NSD-task categorical analyses. This involved using the person and place selective clusters generated from the category localizer task as masks to evaluate the person/place selectivity from the NSD-task selectivity analysis. To ensure the robustness of results, this process was repeated across each data-fold of the NSD-recognition data, such that z-score categorical selectivity maps were generated once from even and once from odd runs. Categorical selectivity scores within category localizer generated clusters did not differ across data-folds (t(22) = 0.049, b = 0.012, 95% CI = [-0.504, 0.528], p = 0.961), confirming the consistency of results, and allowing for scores to be averaged across folds for further analysis. Significant selectivity in the same direction as the category localizer clusters was next evaluated as evidence that the NSD-task selectivity analysis was capturing similar elements of categorically biased responses during recognition memory performance.

Next, we sought to compare how regions demonstrating categorical selectivity related to networks of regions implicated in either recognition or semantic content. To do so, the person and place clusters from the NSD-task were used as masks for generating mean responses from the recognition and semantic content analyses. Again here, we used cross-validation to generate clusters from one half of the data (e.g. even runs) and generated z-scores from the held-out set (e.g. odd runs). Responses were averaged across data-folds and then were compared using mixed effects linear regression, including random subject intercepts and fixed effects of contrast type (recognition/semantic) and cluster type (person/place). As an additional means of measuring overlap between functional regions of the MPC, we quantified the percentage of categorically selective MPC vertices that were also identified as belonging to recognition or semantic clusters. For one subject (S6) no person clusters survived thresholding, so they were excluded from comparisons between person and place clusters.

### Functional beta-series correlation analysis

We also sought to test whether MPC regions involved in recognition decisions or semantic content may be differentially linked to anterior versus posterior hippocampal subregions. We used functional connectivity between hippocampal and MPC subregions to evaluate this relationship. Individual-specific anterior and posterior hippocampal ROIs were designated by manually dividing hippocampi at the uncal apex^123^. Next, task based functional connectivity analyses were performed in a cross-validated manner, where sessions were split into even and odd runs and analyses and recognition or semantic clusters generated from once set of runs were used as masks for comparing connectivity during the held out set of runs.

Task-based MPC to hippocampal subregion connectivity bias was quantified by looking at differences in timeseries correlations between MPC and anterior versus posterior hippocampal voxels during NSD recognition task performance. The normalized volumetric data (z-scored 1.0mm, beta version-2) was first correlated between all MPC and hippocampal voxels separately within each subject and session. For each session, these MPC-hippocampus correlation maps were fisher transformed, and then the average MPC correlation with the anterior hippocampus and within the posterior hippocampus was calculated. Lastly, MPC-hippocampus bias maps for each session were created by subtracting the posterior average from the anterior average for each MPC voxel. This MPC-hippocampal bias score was computed separately for voxels residing in each hemisphere (e.g. left MPC to left hippocampus, left MPC to right hippocampus, etc.). Hippocampal bias scores were input into a second level across session t-test, which determined whether MPC voxels demonstrated a systematic anterior or posterior connectivity bias during task performance. MPC-hippocampal bias reliability maps (t-scores) were then converted to normalized z-scores and projected to three surface layers using nearest-neighbor interpolation. The average across the three surface layers was taken as the final surface bias value. In order to test whether areas responsive to recognition demonstrated different patterns of connectivity to the hippocampus from regions implicated in the semantic content analysis, clusters from those analyses were used as masks. Bias scores were similar across hemispheres and as such combined for statistical comparisons and visualization. First consistency across even and odd runs was tested using mixed effects linear regression, and then differences in hippocampal connectivity bias were examined across recognition and semantic clusters.

### Resting state connectivity analysis

Resting state functional connectivity was measured using the CONN toolbox 21.a^124^ in MATLAB 9.11 and using SPM12^125^ functions. The raw timeseries from the first four resting state sessions (from experiment sessions 21 and 22) for each subject were included. Basic preprocessing performed within the CONN toolbox included motion realignment and unwarping, outlier detection for later scrubbing, direct rigid registration to the 1.0mm T1 structural. Smoothing (3mm) was applied to make file sizes more manageable for analysis. Next, denoising was performed via linear regression, removing noise components identified from white matter and CSF timeseries, as well as noise stemming from estimated subject motion. Scrubbing was also applied to limit the impact of outlier TRs, identified during preprocessing. Lastly, recognition and semantic content cluster masks from across hemispheres were used as seed regions. Seed-to-voxel correlation maps were computed as the Fisher-transformed bivariate correlation coefficients between the mean ROI timeseries and each individual voxel timeseries.

Next, resting state correlation bias scores were calculated in a similar manner as the task-based correlation bias scores above. Recognition cluster correlations were averaged separately for all anterior and posterior hippocampal voxels, and the difference between the average anterior minus posterior voxels was taken as the hippocampal-bias score. The same was done for the semantic clusters. Scores across hemispheres were similar, so were collapsed for subsequent analyses. Cluster-extracted hippocampal-bias scores were entered into a linear mixed effects regression to evaluate the relationship between clusters and hippocampal connectivity during rest.

### Image Permissions

All images used in this manuscript and experiment were from publicly available online resources (COCO stimulus set: https://cocodataset.org/#home; category localizer: https://github.com/VPNL/fLoc#image-sets), and their use here does not conflict with copyright policies therein. Specific images used in this manuscript and their corresponding creative commons licenses can be found for Figure 1: https://www.flickr.com/photos/christophercraig/5580617778/, https://www.flickr.com/photos/scottsherrin/3857178115/ and for Figure 4: https://www.flickr.com/photos/waterarchives/5629839290/, https://www.flickr.com/photos/41809202@N00/4512212885/, https://www.flickr.com/photos/lizacherian/5686169973/, https://github.com/VPNL/fLoc.

## Acknowledgments

This work was supported by NIH grants F32MH130027 to S.R.K and R01MH129439 to B.L.F.

